# Circulating exosomes from Alzheimer’s disease suppress VE-cadherin expression and induce barrier dysfunction in recipient brain microvascular endothelial cell

**DOI:** 10.1101/2023.04.03.535441

**Authors:** Jiani Bei, Ernesto G. Miranda-Morales, Qini Gan, Yuan Qiu, Sorosh Husseinzadeh, Jia Yi Liew, Qing Chang, Balaji Krishnan, Angelo Gaitas, Subo Yuan, Michelle Felicella, Wei Qiao Qiu, Xiang Fang, Bin Gong

## Abstract

**Background:** Blood-brain barrier (BBB) breakdown is a component of the progression and pathology of Alzheimer’s disease (AD). BBB dysfunction is primarily caused by reduced or disorganized tight junction or adherens junction proteins of brain microvascular endothelial cell (BMEC). While there is growing evidence of tight junction disruption in BMECs in AD, the functional role of adherens junctions during BBB dysfunction in AD remains unknown. Exosomes secreted from senescent cells have unique characteristics and contribute to modulating the phenotype of recipient cells. However, it remains unknown if and how these exosomes cause BMEC dysfunction in AD.

**Objectives:** This study aimed to investigate the potential roles of AD circulating exosomes and their RNA cargos in brain endothelial dysfunction in AD.

**Methods:** We isolated exosomes from sera of five cases of AD compared with age- and sex-matched cognitively normal controls using size-exclusion chromatography technology. We validated the qualities and particle sizes of isolated exosomes with nanoparticle tracking analysis and atomic force microscopy. We measured the biomechanical natures of the endothelial barrier of BMECs, the lateral binding forces between live BMECs, using fluidic force miscopy. We visualized the paracellular expressions of the key adherens junction protein VE-cadherin in BMEC cultures and a 3D BBB model that employs primary human BMECs and pericytes with immunostaining and evaluated them using confocal microscopy. We also examined the VE-cadherin signal in brain tissues from five cases of AD and five age- and sex-matched cognitively normal controls.

**Results:** We found that circulating exosomes from AD patients suppress the paracellular expression levels of VE-cadherin and impair the barrier function of recipient BMECs. Immunostaining analysis showed that AD circulating exosomes damage VE-cadherin integrity in a 3D model of microvascular tubule formation. We found that circulating exosomes in AD weaken the BBB depending on the RNA cargos. In parallel, we observed that microvascular VE-cadherin expression is diminished in AD brains compared to normal controls.

**Conclusion:** Using *in vitro* and *ex vivo* models, our study illustrates that circulating exosomes from AD patients play a significant role in mediating the damage effect on adherens junction of recipient BMEC of the BBB in an exosomal RNA-dependent manner. This suggests a novel mechanism of peripheral senescent exosomes for AD risk.

## Introduction

Alzheimer’s disease (AD) is a devastating and ultimately fatal form of neurodegeneration that results in the progressive loss of behavioral and cognitive functions. This neurological disorder imposes a significant emotional and financial burden on families, caregivers, and the healthcare system. However, the mechanism underlying the pathogenesis of AD remains unclear, and there is currently no cure for this disease^1-14^.

Dysregulation of the neurovascular unit and dysfunction of blood–brain barrier (BBB) are critical pathophysiological events in neurodegenerative diseases, including AD^15,16^. The neurovascular unit is a complex functional and anatomical structure composed of cellular and extracellular components that are intimately linked and involved in regulating BBB permeability, cerebral blood flow, and neuronal metabolic activity^15,17,18^.

The BBB is formed by a tightly sealed monolayer of brain microvascular endothelial cells (BMECs), endothelial tight junctions (TJs) and adherens junctions (AJs), working in concert with pericytes to maintain a homeostatic microenvironment for neuronal function^19-25^. BMECs are highly specialized cells with complex TJs and low expression of immune cell adhesion molecules^26^. The regulatory role of the crosstalk between TJs and AJs, in particular the key AJ VE-cadherin (VE-cad)^27^, in stabilizing the BBB has been documented^27-30^.

Loss of some, or most, of the BBB properties is a documented component of the progression and pathology of AD^15,31-34^. BBB dysfunction is primarily caused by reduced or disorganized TJ/AJ proteins or enhanced transendothelial bulk flow of specific molecules by transcytosis^1,2,35^. In early-stage AD patients receiving amyloid-modifying therapies, **a**myloid related imaging abnormalities (ARIA)-including vasogenic edema and microhemorrhage-have been observed^36^, both of which are associated with BBB dysfunction^37^. Evidence from neuropathological, neuroimaging, and cerebrospinal fluid biomarker studies suggests that BBB breakdown is an early biomarker of cognitive dysfunction in human^38^. More than 20 independent postmortem human studies have documented BBB breakdown in AD, revealing BMEC degeneration, TJ disruption, microhemorrhages, and hemosiderin deposits^39^. Although VE-cad in cerebrospinal fluid (CSF) has been reported as a biomarker of endothelial injury in preclinical AD^33^, there is a lack of information regarding the functional role of AJs during BBB dysfunction in AD.

It has been reported that the secretion of inflammatory mediators in senescent cells, a phenomenon known as senescence-associated secretory phenotype (SASP)^40-42^, is activated to affect bystander cells^40,41,43^, contributing to the risk of AD associated with peripheral chronic inflammation^44^. The impacts of SASP are exerted in autocrine and paracrine manners^45,46^. Prolonged exposure to SASP leads to tissue and organ impairment that contributes to the neurovascular degeneration in AD^47-49^. The paracrine SASP mediators include cytokines, chemokines, and the recently recognized exosome^47,50^. Increasing evidence suggests that exosomes secreted from senescent cells have unique characteristics and contribute to modulating the phenotype of recipient cells, in part similar to other soluble SASP factors^46^. Thus, the exosomes secreted from senescent cells, referred to senescence-associated exosomes, appear to be a novel SASP factor^46^.

Extracellular vesicles (EVs) are broadly classified into two main categories, exosomes (also known as small EVs, 50-150 nm) and microvesicles (100-1000 nm), which are distinguished by their cell membrane of origin^51^. Exosome biogenesis begins with the formation of intraluminal vesicles, the intracellular precursors of exosomes, after the inward budding of the membranes of late endosomes^51,52^. These intraluminal vesicles are internalized into a multivesicular body, which transits towards and fuses with the plasma membrane, releasing intraluminal vesicles into the extracellular environment as exosomes^51-53^. Microvesicles are rapidly generated at the plasma membrane by outward budding^51-54^. EVs contain many types of biomolecules and convey signals to a large repertoire of recipient cells, either locally or remotely by ferrying functional cargos, thus contributing to disease pathogenesis^55-62^. There has been increased interest in studying the potential of EVs in AD^63-65^, mainly focusing on their protein contents^66-71^. New evidence^72^ suggests that cultured AD astrocyte-derived EVs induced neuroglial cytotoxicity and endothelial cells (ECs) disruption. Aharon *et al*. reported several overexpressed microRNAs in AD blood-derived mixed EVs^63^. Mounting evidence suggests that many, if not all, of the effects of exosomes on the recipient cell are mediated by exosomal small RNA cargos^73-84^. However, the functional role of RNA cargos in EVs during AD is still unknown.

Given recent evidence indicating that senescence affects exosomes rather than other forms of EVs^85^, in the present study, we isolated circulating exosomes from patient sera and demonstrated that they weaken lateral binding forces (LBFs) between live BMECs, which directly measure the biomechanical nature of the endothelial barrier. This effect was associated with the suppression of paracellular VE-cad in *in vitro* and *ex vivo* models of BBB and was dependent on the exosomal RNA cargos. Furthermore, we observed diminished microvascular VE-cad immunofluorescent (IF) signals in frontal cortex areas of sporadic AD patients compared to cognitively normal controls.

## Results

### 1. AD circulating exosomes (AD-Exos) induce barrier dysfunction in normal recipient BMECs in a dose-dependent manner

BMEC activation or injury is an early and prevalent event in AD^15-18,33,37,38,86-91^. To assess the potential effects of circulating exosomes on BMECs, we isolated exosomes from 0.2 µm-filtered sera of 5 cases of AD with clinical dementia rating (CDR) scores of 3 compared with 5 cases age- and sex-matched cognitively normal controls with CDR 0 (see **Table**), using size-exclusion chromatography (SEC) that was documented with improved integrity, yield, and no aggregation compared with other methods^53,92-95^. We validated exosome qualities and particle sizes (50-150 nm) using EV-specific assays^96,97^ and confirmed that the particles are exclusively distinct from synaptosome-sized particles (0.6 µm to 1.6 µm^98^) (**Fig. 1A-C**) and exosome size distribution and morphology were not significantly altered between groups.

**Table:** Information of control- and AD-samples.

| **Table: Information of control- and AD-samples** |  |  |
| --- | --- | --- |
| Sera for exosome isolation |  |  |
| No. of cases | Cognitively normal (CDR=0) | AD (CDR=3) |
| Gender (M/F) | 2/3 | 2/3 |
| (year) | 62-79 | 62-84 |
| Archival brain tissues (from UTMB and BU) |  |  |
| No. of cases | Cognitively normal (CDR=0) | AD (CDR=3) |
| Gender (M/F) | 3/2 | 3/2 |
| (year) | 65-96 | 66-92 |

**Figure 1.**
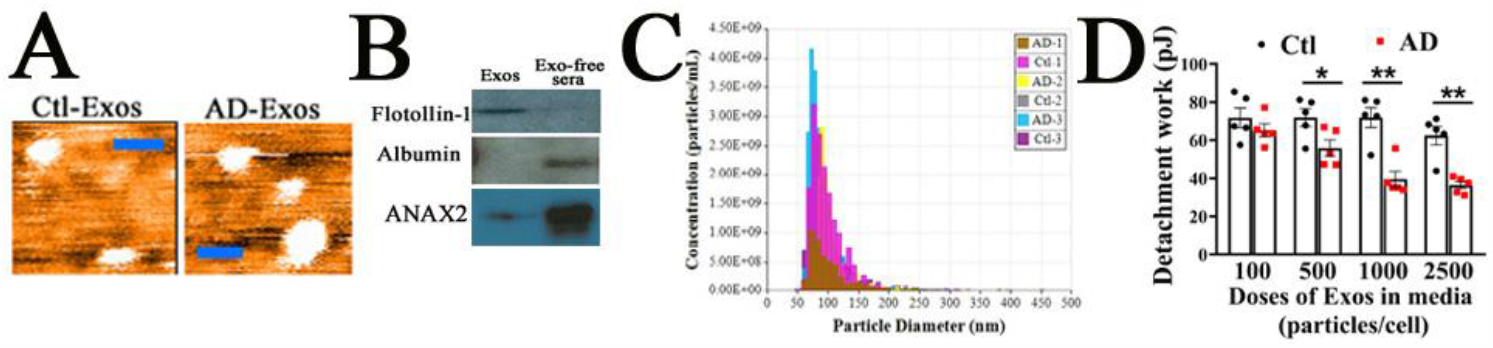
AD-Exos induce barrier dysfunction in normal recipient BMECs in a dose-dependent manner. **(A)** Ctl-Exo and AD-Exo morphologies were verified using atomic force microscopy (AFM) (scale bars, 200 nm). (**B**) Expressions of indicated protein markers in 100 μg proteins of Exos were examined using western immunoblotting. (**C**) The vesicle size distribution of isolated Exos was analyzed using nanoparticle tracking analysis (NTA). (**D**) The lateral binding forces (LBFs) were assessed by measuring the detachment works of live recipient BMECs using fluidic AFM 72 hr after exposure to AD-Exos at 100, 500, 1000, and 2500 particles/cell. *, p < 0.05; **, p < 0.01 (Determined by one-way ANOVA).

The intercellular multi-protein junctional complexes in ECs play a fundamental role in sealing the lateral space between cells against unbinding forces at the lateral contact sites^99^. Therefore, the measurement of lateral binding forces (LBFs) between BMECs is crucial in evaluating the precise biomechanical features underlying BBB dysfunctions. To directly measure the LBFs between live BMECs Employing, a new biomechanical strategy using fluidic force microscopy (fluidic AFM)^97,100,101^ was employed (. The results revealed that the LBFs between live BMECs were reduced in a dose-dependent manner after 72-hr treatment with AD-Exos (**Fig. 1D**, n=5/group) (**Suppl Fig. 1**), suggesting AD-Exos induce barrier dysfunction in normal recipient BMECs.

### 2. AD-Exos reduce paracellular expression of VE-cad in a manner dependent on exosomal RNA cargos in BMEC cultures

During IF assays of BMECs, we observed marked suppression of the paracellular signal of VE-cad in recipient BMEC 72 hr post exposure to AD-Exos, but not control exosomes (Ctl-Exos) (1000 particle/cell in media) (**Fig. 2A**).

**Figure 2.**
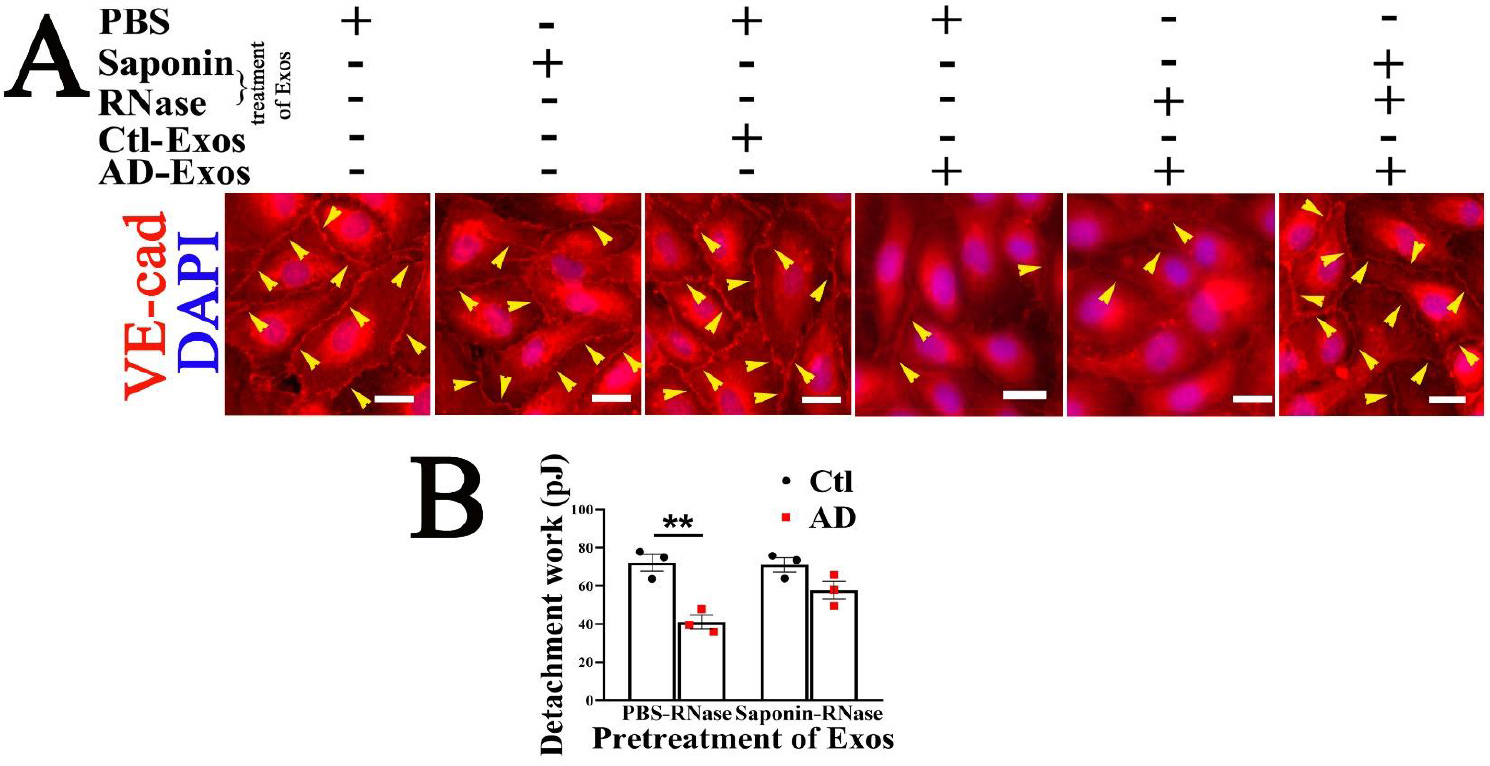
AD-Exos reduce paracellular expression of VE-cad (arrow heads in **A**) and weaken the barrier function of recipient BMECs in a manner dependent of exosomal RNA cargos. (**A**) Representative IF staining of VE-cad in BMECs in BMEC cultures 72 hr after exposure to Ctl-Exos or AD-Exos (at a concentration of 1000 particles/cell) that were pretreated with 20 µg/mL ribonuclease (RNase) in the presence or absence of 0.1% saponin as previously reported^97,101^. Scale bars, 20 µm. (**B**) The LBFs were assessed by measuring the detachment works of live recipient BMECs using fluidic AFM 72 hr after exposure to AD-Exos (at 1000 particles/cell) that were pretreated with 20 µg/mL ribonuclease (RNase) in the presence or absence of 0.1% saponin. *, p < 0.05; **, p < 0.01 (Determined by one-way ANOVA).

Exosome can convey signals to a large repertoire of neighboring and distant recipient cells by ferrying functional cargos. Although exosomes contain both proteins and RNAs, many, if not all, of the effects of Exos on the recipient cell are mediated by exosomal RNA cargos^73-79^. To investigate the role of exosomal RNA cargos during AD-Exo-induced suppression of VE-cad, we pretreated exosomal cargos with RNase using our reported saponin-assisted active permeabilization method^96^. This pretreatment mitigates the detrimental effects of AD-Exos on BMEC paracellular expression of VE-cad (**Fig. 2A**), suggesting that AD-Exos reduce paracellular expressions of VE-cad depending on exosomal RNA cargos.

To confirm the effect of pretreatment of AD-Exos on the recipient BMEC barrier function, we further measured the LBFs of live BMECs following exposure to different pretreated-AD-Exos prior to applying in culture media. Information from LBF assays confirmed that the detrimental effects of AD-Exos on recipient BMEC barrier functions depend on exosomal RNA cargos (**Fig. 2B**, n=3/group).

### 3. AD-Exos perturb the integrity of VE-cad in 3D BBB models

Using another model for the BBB, 3D collagen scaffolds of microvascular constructs have been utilized to replicate *in vivo* cell-to-cell interfaces during microvascular tubule formation^31^. Our 3D BBB models employed primary human BMECs and pericytes (**Fig. 3A** and **B**). Again, confocal image analysis of IF staining revealed a reduction in the coverage of microvasculature VE-cad, which is a targeted measure of BBB structures^31,32,102^, in AD-Exos-treated constructs compared to the Ctl-Exo-treated group 72 hr after exposure to different exosomes in media (**Fig. 3C** and **D**, n=25/group). This further suggests that AD-Exos perturb the integrity of VE-cadherin in BMECs of the BBB.

**Figure 3.**
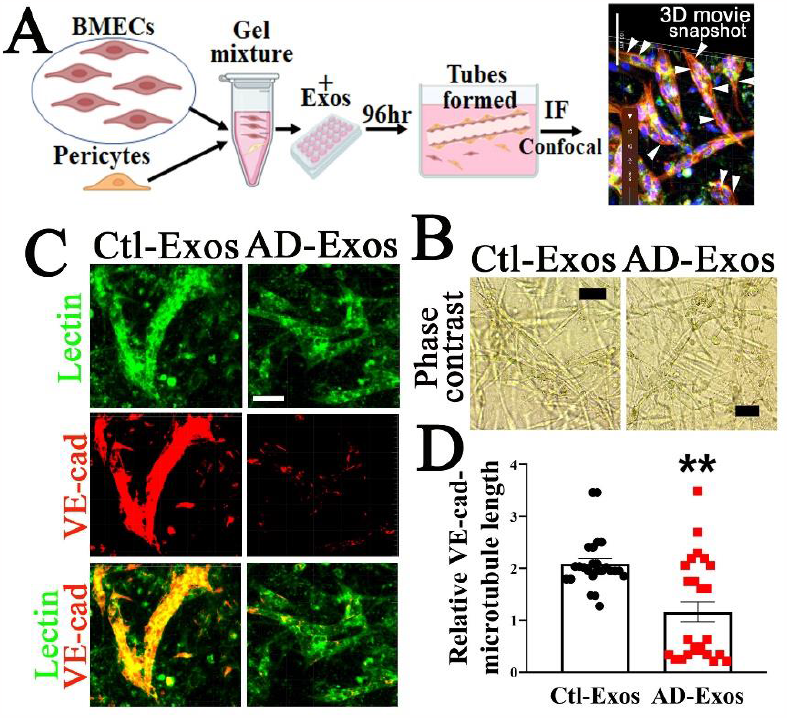
AD-Exos disrupt the integrity of VE-cad in 3D BBB models. (**A**) The experiments used human primary BMEC-pericyte collagen scaffold tubule formation systems. A schematic representation of the experiments is shown. (**B**) Representative phase contrast images of the 3D constructs at 72 hr post-exposure to Ctl-Exos and AD-Exos at 2500 particles/cell are shown. Scale bars, 50 µm. Representative confocal images (**C**) and relative VE-cad-labeled microvascular tubule lengths (**D**) in Ctl-Exos- vs. AD-Exos-treated (at 2500 particles/cell) 3D contracts at 72 hr post-exposure are presented. Scale bars, 50 µm. **, p < 0.01 (Determined by *t*-test).

### 4. In AD brains, VE-cad expression is diminished

VE-cad is a key protein of AJs in EC^103^. In parallel, we examine the IF signal of VE-cad in brain tissues from 5 cases (see **Table**) of AD with CDR scores of 3 compared with 5 cases age- and sex-matched cognitively normal controls with CDR 0. Indeed, we observed that in all AD brains, microvascular VE-cad IF signals in frontal cortex areas were diminished compared to all controls (**Fig. 4**). In contrast,, the expression of VE-cad in lung tissues from the same participants did not show a difference between the 3 cases each of the AD and control groups (**Suppl Fig. 2**), suggesting the detrimental effects of AD-Exos may be cell-type dependent.

**Figure 4.**
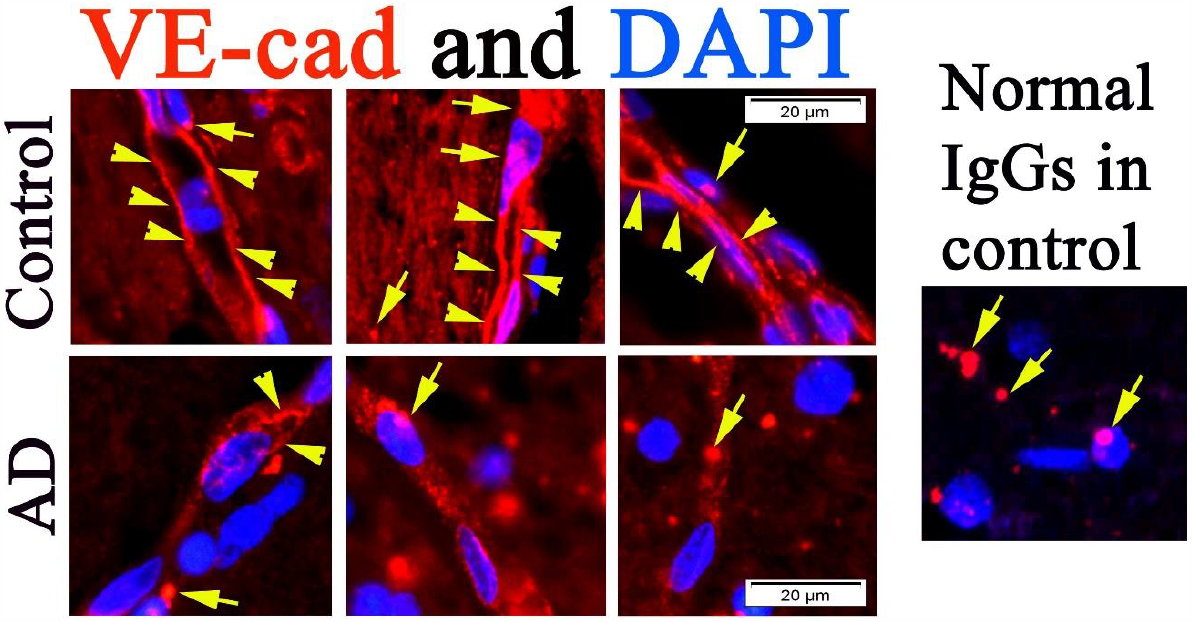
Representative immunofluorescence staining of VE-cad in brain frontal cortex areas from individuals with AD who had a CDR score of 3 (n=5) and age- and sex-matched cognitively normal controls with CDR scores of 0 (n=5). VE-cad signals (indicated by arrow heads) were visualized using rabbit anti-human VE-cad antibodies and nuclei were counterstained with DAPI (blue). Normal rabbit IgGs were utilized as a negative control to demonstrate the background fluorescence (indicated by arrows) in formalin-fixed paraffin-embedded human brain tissue^144^. Scale bars, 20 µm.

## Discussion

In the present study, evidence from immunostaining analysis and nanomechanical assays suggested that AD circulating exosome can suppress paracellular expression levels of VE-cad and impair the barrier function of recipient BMECs. Our 3D model of microvascular tubule formation also demonstrated the damaging effects of AD-Exos on VE-cad integrity. Additionally, we found that AD-Exos weaken the BBB depending on the RNA cargo carried by exosomes. This was demonstrated by using saponin-assisted active permeabilization of exosomes before applications in BMEC culture and 3D BBB models. Furthermore, we observed that microvascular VE-cad signals diminished in five AD cases with CDR 3.

AD is the most common type of dementia accounting for an estimated 60% to 80% of cases^104^. Pathologically, the most documented attributes of AD are the build-up of beta-amyloid plaques on the outside of neurons and the neurofibrillary tangles (NFT) made up of hyperphosphorylated tau (P-tau) and found intraneurally^105^. Clinically, patients demonstrate a loss in cognitive function, including memory, language and visuospatial alterations. Additionally, behavioral disorders such as depression or apathy may also be present^106^. Some genetic associations have been described in AD including those for early AD (under 65 years) and late AD (after 65 years). The latter is more common and usually occurs due to a polymorphism in the *APOE* gene. Whereas the former has been linked with mutations in the genes *APP, PSEN1* and *PSEN2* (amyloid precursor protein, presenilin 1 and presenilin 2, respectively)^107^. Given the increase in the aging population worldwide, evidence indicates that dementia has become an economic burden worldwide. According to the Delphi Consensus Study, a worldwide estimate for the number of vascular dementia cases could reach up to 81.1 million by 2040^108^. Therefore, improved understanding of the proposed pathophysiological mechanisms of AD is vital in order to improve diagnosis and in the discovery of novel therapies. This would allow worldwide authorities to better improve the current socioeconomic burden caused by AD.

Microvasculature injury is a common observation in most AD brains, even in preclinical stages^109,110^. While VE-cad in cerebrospinal fluid (CSF) was reported as a biomarker of endothelial injury in preclinical AD^33^, there is a lack of information on the functional role of AJs during BBB dysfunction in AD. BBB properties are primarily determined by the AJs and TJs^26,111^, with BMECs forming extremely tight TJs that are distinct from endothelial or epithelial TJs elsewhere, which have fenestrations and high rate of pinocytosis^112,113^. Disruption of the TJs at the BBB is a common occurrence in many neurological disorders^114-119^. TJs are located at the most apical position among all junctional components and are core structures that help to seal the BMECs at the BBB^114,120^. Studies on developmental stage suggest a high level of interdependency between AJs and TJs during the formation of cell–cell contacts between ECs^114,120-122^. Interestingly, the expression level of VE-cad in CSF is increased in preclinical AD and correlates with cognitive impairment^33^, but underlying mechanisms remain unknown. We noted that in AD brains, microvascular endothelial AJ protein VE-cad was downregulated. However, the potential interplay between AJs and TJs during the pathogenesis of BBB dysfunction in AD remains to be addressed in our future studies.

In the present study, we revealed that AD-Exos induce detrimental effects on recipient BMECs that depend on their RNA cargos. Despite the abundant presence of RNases in blood, extracellular RNAs have been discovered, leading to the proposal of a scenario in which RNAs are encapsulated in exosomes^123-126^. Exosomal RNA cargo mostly consists of small noncoding RNAs (sncRNAs), mainly tRNA fragments^80,81^ or microRNAs^74-77,127-129^, and mounting evidence indicates that many effects of exosomes are mediated by them^74-77,80,81,127-129^. Our findings suggest that AD-Exos carry specific RNA cargos that can cause damage to BMEC barriers, which are critical for maintaining the integrity of the BBB. Further studies are needed to identify and characterize the specific RNA cargos responsible for these harmful effects and elucidate the mechanism underlying the biogenesis of these specific RNA cargos.

The correlation between endothelial inflammation/activation and neurovascular degeneration is providing new insights into the pathogenesis of AD^32,44^. Microvascular endothelial dysfunction refers to impaired functioning of the lining of capillary blood vessels and is characterized by endothelial surface proadhesive activation, enhanced coagulation, microthrombosis, and endothelial barrier breakdown^130^. Additionally, growing evidence supports the essential role of pericytes in maintaining a healthy and optimally functioning BBB^31,131^. Our recent work, which utilized BMEC culture and 3D microvascular tubule construct, revealed the harmful effects of AD-Exo on recipient endothelial barrier function in association with VE-cad. However, the potential role(s) of pericytes in this process remain to be identified. To better understand the underlying mechanism(s) of BBB dysfunction in AD, it is necessary to evaluate the complex interactions between the various components of the neurovascular unit, including BMECs, pericytes, astrocytes, and neurons^15,132,133^, *in vivo* in future studies.

Our findings suggest that circulating exosomes in individuals with AD can inhibit the expression of VE-cad, an AJ molecule that is visualized to be decreased in AD brains, leading to barrier dysfunction in BMECs. This disruption of the BBB may contribute to the development of AD by facilitating the infiltration of harmful molecules and cells into the central nervous system. Our study highlights the importance of considering the role of exosomes in the pathogenesis of AD and provides a potential mechanism by which these exosomes contribute to disease progression. Further investigations are necessary to identify the specific RNA cargos responsible for these effects and to develop targeted therapies aimed at mitigating the detrimental effects of AD-Exos on BMECs and other neurovascular unit cells.

## Materials and Methods

### Human sample selection

The Collaborative AD Program at UTMB data collection procedures were approved by the UTMB Institutional Review Board. Participants (or their Legally Authorized Representatives) provided written informed consent to participate in the study. The study adhered to the most recent guidelines set by The World Medical Association Declaration of Helsinki set by the 64^th^ WMA General Assembly. Upon a patient’s scheduled visit, the clinical diagnosis of AD dementia was confirmed by a board-certified neurologist. Imaging studies such as functional MRI and positron emission tomography in conjunction with cognitive tests such as the Montreal Cognitive Assessment and Mini Mental State Exam were utilized for diagnostic purposes and to determine the stage of the AD dementia. The Clinical Dementia Rating (CDR) Scale, a global clinical assessment based on direct patient and caregiver information, was also used for determining dementia severity. CDR is based on a scale of 5 points and characterized by six domains of cognitive and functional performance including community affairs, home & hobbies, judgment & problem solving, memory, orientation, and personal care. Scoring is calculated through an algorithm. Once determined, the 5 possible points are: 0 = Normal, 0.5 = Very Mild Dementia, 1 = Mild Dementia, 2 = Moderate Dementia, and 3 = Severe Dementia. MMSE was also used to determine severity of dementia: Mild AD = MMSE 21–26 points. Moderate AD = MMSE 15–20 points. Moderately severe/severe AD = MMSE<15 points. For proper identification of the blood samples, a standardized labeling system was implemented, while safeguarding the anonymity of the participants. Samples were transported following standardized guidelines and processed in the laboratory.

Procurement of human post-mortem formalin-fixed brain tissues for research were approved by the University of Texas Medical Branch (UTMB; Galveston, TX), Committee on Research with Post-Mortem Specimens. All appropriate post-mortem permissions, precautions, and procedures done were according to the guidelines of the College of American Pathologists^134^. Informed consent was obtained from all study participants, and the study protocol was approved by the Institutional Review Board of Boston University Medical Campus.

Informed consent was obtained from all study participants, and the study protocol was approved by the Institutional Review Board of Boston University Medical Campus.

### Exo-isolation^96^

The serum sample (200 µl) was passed through 0.2µm syringe filters first. Following the manufacturer’s instructions, filtered plasma was placed onto the qEVoriginal column (Izon, New Zealand) for SEC isolation. The number 7 to 9 fractions were collected as the Exo-enriched fractions, which were concentrated using 100,000 MWCO PES Vivaspin centrifugal filters (Thermo Fisher Scientific). Exo samples (in 200 µl PBS) were stored at -80°C prior to use in downstream assays. Following the manufacturer’s instructions, three fractions (the number 13 to 15 fractions) from the qEVoriginal column isolation were collected as the Exo-free, high-protein plasma fractions, which were concentrated using centrifugal filters and used in western immunoblotting as a control.

For saponin-assisted active exosomal permeabilization pretreatment of Exos using RNase as reported^96^, Exo samples (1 × 10^9^ particles/mL) and RNase (20 µg/mL) (Thermo Fisher Scientific) were incubated with 0.1 mg/ml saponin (Thermo Fisher Scientific) at room temperature for 15 min. After rinsing using PBS, Nanoparticle tracking analysis (NTA) was performed using a qNano Gold system (Izon, New Zealand) to determine the size and concentration of extracellular vesicle particles of each Exo sample. Exo samples were concentrated using 100,000 MWCO PES Vivaspin centrifugal filters.

### Nanoparticle tracking analysis (NTA)^97^

NTA was performed using the TRPS technique and analyzed on a qNano Gold system (Izon, Medford, MA) to determine the size and concentration of EV particles. With the qNano instrument, an electric current between the two fluid chambers is disrupted when a particle passes through a nanopore NP150 with an analysis range of 70 nm to 420 nm, causing a blockade event to be recorded. The magnitude of the event is proportional to the number of particles traversing the pore, and the blockade rate directly relates to particle concentration that is measured particle by particle. The results can be calibrated using a single-point calibration under the same measurement conditions used for EV particles (stretch, voltage, and pressure)^135^. CPC200 calibration particles (Izon) were diluted in filtered Izon-supplied electrolyte at 1:500 to equilibrate the system prior to measuring EVs and for use as a reference. Isolated extracellular vesicle samples were diluted at 0, 1:10, 1:100, and 1:1,000. For the measurements, a 35 µl sample was added to the upper fluid chamber and two working pressures, 5 mbar and 10 mbar, were applied under a current of 120 nA. Particle rates between 200 and 1500 particles/min were obtained. The size, density, and distribution of the particles were analyzed using qNano software (Izon).

### Imaging of label-free extracellular vesicles using atomic force microscopy (AFM) ^97^

To obtain height images, the purified and concentrated EV sample was diluted at 1:50, 1:500, and 1:1000 with molecular grade water. Glass coverslips were cleaned with ethanol and acetone before it was coated with the diluted exosomes, before being examined using an AFM (Nanosurf AG, Liestal, Switzerland) using contact mode in the air. A PPP-FMR-50 probe (0.5-9.5N/m, 225µm in length and 28µm in width, Nanosensors) was used. The parameters of the cantilever were calibrated using the Sader *et al*. method^136^. The cantilever was approached to the sample under the setpoint of 20 nN, and topography scanning was done using the following parameters: 256 points per line, 1.5 seconds per line in a 5-µm x 5-µm image.

### BMEC culture

The BMEC culture was established in a similar way to what was previously described^97^. Primary human BMECs were cultured in Endothelial Cell Growth Medium (Cell Applications). Treatments were started 24 hr after seeding and lasted until designed time for downstream assays.

### Immunofluorescent (IF) staining

Unless otherwise indicated, all IF reagents were obtained from Thermo Fisher Scientific. For IF of VE-cad in BMEC cultures, fixed BMECs were incubated with anti-VE-cad rabbit antibody (ABclonal) (1:100). A rabbit normal IgG served as a negative control. Nuclei were counterstained with 4′,6-diamidino-2-phenylindole (DAPI). For the IF of VE-cad in brain tissue sections, slides were incubated with anti-VE-cad rabbit antibody (1:100) overnight at 4°C, after antigen retrieval and blocking. Negative controls were incubated with normal rabbit IgGs. Nuclei were stained with DAPI.

### BMEC-pericyte-collagen matrix 3D-scaffold BBB model

The 3D collagen scaffolds of microvascular tubule formation model were prepared using 3D Tubule Formation kit (Catalog 8698, ScienCell) as previously described^31,137^. Briefly, collagen solution was mixed with cell culture medium containing BMECs and pericytes at a ratio of 5:1, and then added into a well of a 24-well plate at 75 µl/well. The mixture was incubated at 37°C for 60 minutes for polymerization before adding complete 3D Medium (ScienCell) supplemented with different exosomes at 2500 particle/cell. The plate was then transferred to a 5% CO2 incubator and cultured at 37°C. The scaffolds were imaged by phase contrast daily to assess the formation of microvascular networks for 5 days before fixation with 4% paraformaldehyde.

IF of VE-cad coupled with lectin staining, which outlines microvascular patterns^102,138^, was performed on fixed scaffolds mounting on slides. The scaffolds were permeabilized in Triton-100 for 20 minutes before incubation with anti-VE-cad rabbit antibody (1:50) overnight at 4°C. After rinsing three times with 0.1% Tween in PBS, the scaffolds were incubated with Alexa Fluor 594-conjugated goat anti-rabbit IgG (Thermo Fisher) (1:1000) overnight at 4°C. A rabbit normal IgG (Thermo Fisher) served as a negative control. After rinsing, the scaffolds were incubated with Lycopersicon Esculentum Lectin-DyLight 488 (Vector Laboratories) (1:100) for 2 hours at room temperature. Nuclei were stained with DAPI. All stained constructs were analyzed using a Nikon A1R MP ECLIPSE T*i* confocal microscope as previously reported^139^. The relative VE-cad-labeled lengths of microvascular tubule formation were quantified using ImageJ software, modified as described^32^. Briefly, five selected arears in the center of the stained constructs were scanned. Z-stacks were collected at 1-µm steps for a total imaging depth of 15 µm. Confocal microscopy-recorded images were processed and exported using Imaris Viewer (Oxford instruments), an open platform supporting confocal 3D volume rendering and quantitative analysis^140^. In this study, VE-cad signals were visualized not only at microvascular tubule structures but also accumulated in non-tubule structure areas, while lectin-labeled signals mainly depicted the tubular structures^102,138^ in the scaffolds. In ImageJ program, to quantify the relative VE-cad-labeled microvascular tubule lengths, all lectin channel (green) images were processed using the ImageJ/Process/Image Calculator/Original-Image/Subtract/FITCγ-reduced image/analyze/histogram/RGB tool to filter out fluorescent signals in non-tubule areas. After setting these lectin-labeled tubule structures as a map, lengths of VE-cad-labeled single microtubules were measured using the ImageJ/Straight/Analyze/Measure tool. The relative length of VE-cad-labeled were expressed as the ratio of the cumulation of VE-cad tubule lengths in each field to cells (DAPI labeled nuclei).

### LBF measurement using Fluidic AFM micropipette

The fluidic AFM micropipette was used to measure the lateral binding force (LBF) between live BMECs and assess endothelial barrier function, as previously described^97,101^. Briefly, the fluidic AFM chamber was filled with medium, and the temperature was maintained at 37°C by the Temperature Controller (NanoSurf). A micropipette with an 8 μm aperture and spring constant of 2 N/m (CytoSurge) was calibrated using the Sader method in air and liquid by performing deflection and crosstalk compensation. Using an inverted microscope, the cantilever was kept at 20 mbar positive pressure by the fluidic pressure controller (Cytosurge) of the FluidFM system and was brought into contact with the surface of the target BMEC. After a set pause of 10 sec, suction pressure of - 200 mbar decreased to -400 mbar to ensure a seal between the apical surface of the cell and the aperture of the hollow cantilever. The cantilever was vertically retracted to a distance of 100 µm to separate the targeted BMEC from the substrate and the monolayer. The unbinding effort was assessed by recording the deflection of the cantilever, which was calculated by integrating the area under the force-distance (F-D) curve, and measuring the work done (in picojoules [pJ]) using software as we described^141,142^. The captured BMEC was released prior to moving the cantilever to another individual BMEC that did not contact other cells in the same culture. The LBF value was obtained using the method of Sancho *et al*.^143^.

### Statistical analysis

All data were presented as mean± standard error of the mean and analyzed using SPSS version 22.0 (IBM^®^ SPSS Statistics, Armonk, NY). A two-tailed Student’s *t*-test or one-way analysis of variance (ANOVA) was used to explore statistical differences. If the ANOVA revealed a significant difference, a post hoc Tukey’s test was further adopted to assess the pairwise comparison between groups. The level of statistical significance for all analyses was set at *p*<0.05.

## Acknowledgments

We gratefully acknowledge Dr. Kimberly Schuenke for her reviews and editing the manuscript. This work was supported by NIH grants R01AI121012 (B.G.), R21AI137785 (B.G.), R21AI154211(B.G.), R21AG066060 (X.F.), R61AG075725 (X.F.), 1RF1AG075832 (W.Q.Q.), and John Sealy Distinguished Chair in Alzheimer’s diseases (XF). The funders had no role in the study design, data collection and analysis, decision to publish, or preparation of the manuscript.

## Authorship Contributions

W.Q.Q., X.F. and B.G. conceptualized the studies. J.B., E.G.M., Q.G., Y.Q., S.H., J.L., and Q.C. conducted the experiments. B.K., A.G., S.Y., M.F, W.Q.Q., X.F., and B.G. designed the studies. J.B., Y.Q., S.H., W.Q.Q., X.F., and B.G. analyzed the data. J.B., Y.Q., Q.C., A.G., S.Y., and B.G. developed the methodology. J.B., E.G.M., Y.Q., W.Q.Q., X.F., and B.G. wrote the manuscript.

